# Hemolytik2: An Updated Database of Hemolytic Peptides and Proteins

**DOI:** 10.1101/2025.05.12.653624

**Authors:** Ayushi Singh, Kavin Raj SA, Anand Singh Rathore, Gajendra P. S. Raghava

**Affiliations:** Department of Computational Biology, Indraprastha Institute of Information Technology, Okhla Phase 3, New Delhi-110020, India

**Keywords:** Hemolytic Activity, Hemotoxin, Chemical Modifications, Physical Modifications, Database

## Abstract

Hemolytik 2.0 (http://webs.iiitd.edu.in/raghava/hemolytik2/) is a comprehensive, manually curated database that provides experimentally validated information on both hemolytic and non-hemolytic peptides. Data were meticulously extracted from peer-reviewed publications and established peptide repositories, including the Antimicrobial Peptide Database (APD), UniProt, the Collection of Anti-Microbial Peptides (CAMP-R4), and the Dragon Antimicrobial Peptide Database (DAMPD). This updated version of the original Hemolytik resource encompasses 13,215 entries, representing approximately 8,700 unique peptides. Each entry in Hemolytik 2.0 offers detailed annotations, including peptide name, amino acid sequence, biological source and origin, functional characterization, terminal modifications, stereochemistry, structural classification (linear or cyclic), and experimentally determined hemolytic activity. In addition, the database provides molecular representations of peptides in SMILES (Simplified Molecular Input Line Entry System) format, alongside predicted tertiary structures and annotated secondary structural states. Additionally, a RESTful API has been integrated into the Hemolytik 2.0 repository to enable programmatic access and automated retrieval of peptide data. Hemolytik 2.0 serves as a valuable resource for the scientific community, particularly for researchers involved in the design and development of therapeutic peptides, by facilitating the identification and optimization of peptide candidates with minimal hemolytic potential and enhanced safety profiles.

## Introduction

Peptides have emerged as powerful therapeutic agents due to their unique ability to modulate a wide range of biological processes with high target specificity, low toxicity, and favorable pharmacokinetic profiles^1,2^. Over the past decade, advances in peptide synthesis and screening technologies have led to the discovery of numerous bioactive peptides with potential applications in oncology, infectious diseases, autoimmune disorders, and metabolic conditions^3–6^. However, despite their therapeutic promise, many peptide candidates exhibit undesirable toxicity, which may be divided into three main categories: immunotoxicity, cytotoxicity, and hemotoxicity. Hemotoxicity which lyses red blood cells^7,8^. i.e., the ability to lyse red blood cells, which poses a serious safety risk and limits their clinical utility. Hemolytic concentration (HC50), which is the concentration at which 50% of normal human erythrocytes lyse under physiological circumstances, is a frequently used measure of peptide toxicity^9^. Peptides exert hemolytic activity primarily by disrupting the integrity of red blood cell (RBC) membranes through mechanisms such as pore formation, lipid displacement, and self-assembly on membrane surfaces^10–12^.

Hemolysis, often resulting from interactions with the phospholipid bilayer of erythrocyte membranes, is a critical parameter in the early stages of peptide drug development. The ability to accurately predict or assess a peptide’s hemolytic potential is, therefore, essential for optimizing lead compounds and minimizing off-target cytotoxic effects^13–15^. Yet, despite its importance, hemolytic toxicity is often overlooked during early-stage screening due to the limited availability of comprehensive and accessible resources. Most existing datasets are either fragmented across various publications or lack the structural and physicochemical annotations necessary for advanced computational analyses.

To address this gap, we present Hemolytik 2.0, an extensively curated and updated version of the Hemolytik database. Hemolytik 2.0 provides an enriched repository of experimentally validated hemolytic and non-hemolytic peptides, integrating information from a wide range of sources including the Antimicrobial Peptide Database (APD)^16^, UniProt^17^, Collection of Anti-Microbial Peptides (CAMP-R4)^18^, and the Dragon Antimicrobial Peptide Database (DAMPD)^19^. The current version encompasses 13,215 entries, representing approximately 8,700 unique peptides, and provides detailed annotations such as peptide names, sequences, origin, known biological functions, structural class (linear or cyclic), stereochemistry, terminal modifications, and hemolytic profiles. Importantly, Hemolytik 2.0 includes SMILES representations and predicted tertiary structures of peptides, along with secondary structural state assignments, offering users a unique opportunity to explore structure-activity relationships. In addition, a RESTful API has been integrated to support automated data access, thereby enhancing the database’s utility for large-scale machine-learning applications and drug design workflows.

We believe Hemolytik 2.0 will serve as a vital platform for researchers in the field of peptide therapeutics, aiding in the design of safer, non-hemolytic analogs, and accelerating the development of clinically viable peptide drugs.

## Methods

### Data Procurement and Compilation

Data were manually curated from various published research literature and databases. To identify relevant published literature, an advanced search was conducted within PubMed using the following query: *“((((hemolysis) OR (hemolytic)) OR (hemotoxin)) AND (peptide) AND (2013:2024[pdat])) NOT ((((hemolysis) OR (hemolytic)) OR (hemotoxin)) AND (peptide) AND ((review[Filter] OR systematic review[Filter]) AND (2013:2024[pdat]))”*. This strategy was specifically designed to include only original research articles published between 2013 and 2024 that experimentally evaluated the hemolytic activity of peptides using hemolysis assays. As a result, a total of 4,533 articles were shortlisted for data extraction. As a result, a total of 4,533 articles were shortlisted for data extraction. Additionally, data were gathered from well-established peptide repositories, including CAMP R4 (35 peptides), DAMPD (6 peptides), APD3 (179 peptides), and UniProt (268 peptides), using relevant keywords such as “hemolysis” and “hemolytic”. For each peptide entry, detailed information was extracted, including the amino acid sequence, terminal modifications, chirality, biological origin, source of red blood cells (RBCs) used in the assay, and experimentally determined hemolytic activity.

### Database architecture and web interface

After the collection and compilation of data, the next step was to construct the database which was completed by using Apache HTTP Server and MySQL. The front-end interface of the database was created using HTML, CSS, PHP, and Javascript. The choice of Apache, MySQL, and PHP was driven by their open-source nature and cross-platform compatibility. PHP and PERL programming languages were used to write all the common gateway interface and database interface scripts. MySQL functions as the backend and serves as an object-related database management system (RDBMS), providing the necessary commands for data retrieval and storage. The architecture of Hemolytik 2.0 is depicted in Figure 1.

**Figure 1.**
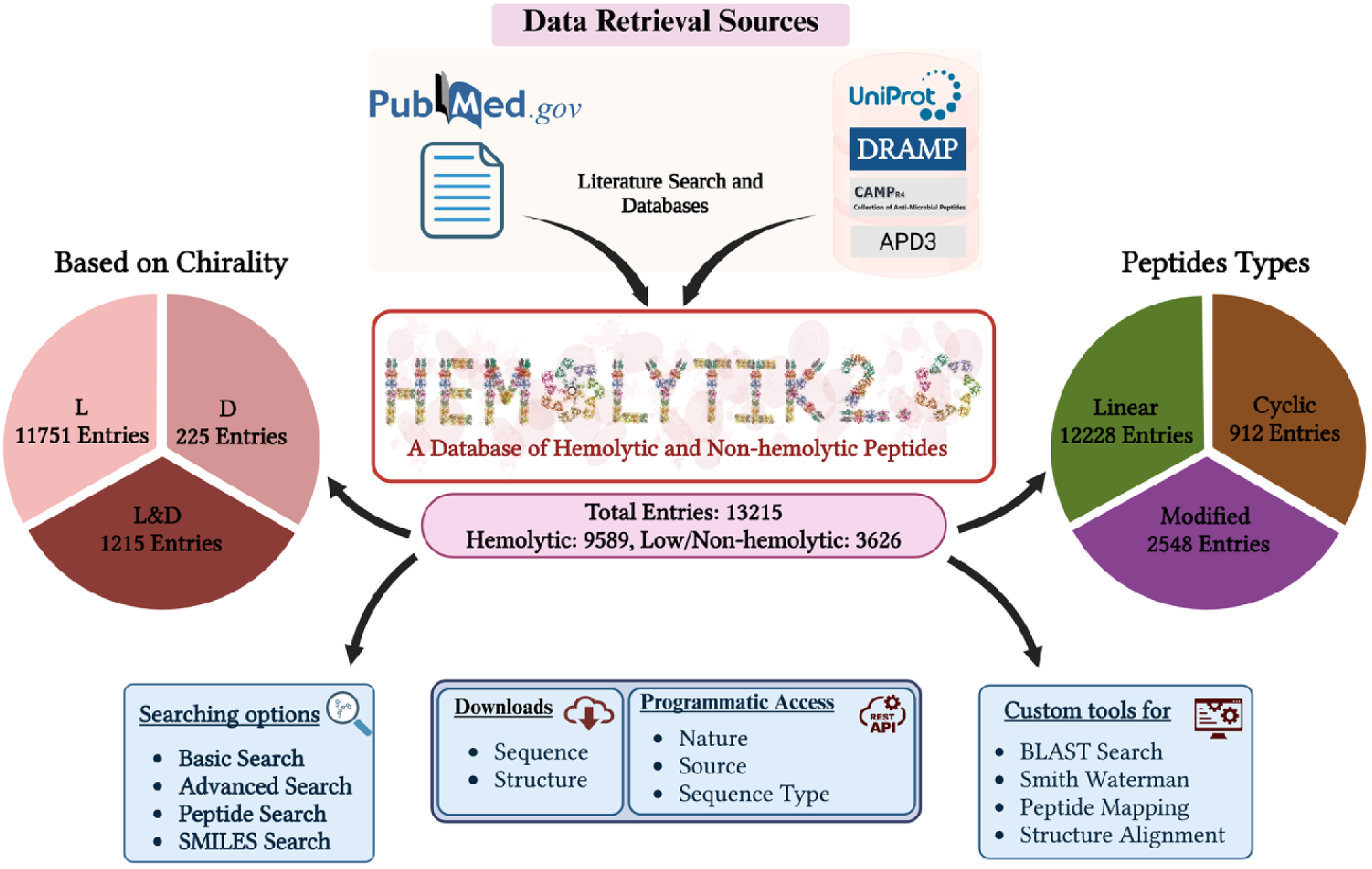
Illustration of the overall architecture of Hemolytik 2.0.

### Data statistics

The updated version of Hemolytik 2.0 includes 13215 entries. The stats are categorized into two parts:

1. **Statistics of all Hemolytic Peptides:** This portion includes stats of all hemolytic peptides. Hemolytic activity was evaluated through assays performed on red blood cells derived from different species. Here, we found various sources of hemolytic activity, i.e. red blood cells isolated such as humans, horses, pigs, porcine, rats, rabbits, fish, etc. Therefore, when we blended information. As a result, we got 20 sources, as shown in Figure 2. Mostly found entries had Human (9255) as source, following horse (1062), sheep (840), mouse (595), etc. their entries were not only single for individual peptides but also multiple entries at different concentrations (mouse and human at 50 µM and 100 µM). The peptides included in the Hemolytik 2.0 database represent various therapeutic categories. The different biological functions represented by the peptide, such as antimicrobial, anticancer, antibacterial, cell-penetrating, etc., most of the entries possess antimicrobial nature (5760) followed by antibacterial (2841). The peptides have been further categorized based on their stereochemistry (L, D or mixed L/D amino acids) and conformation (Linear, Cyclic). There are a total of 12,211 entries for linear peptides, 906 for cyclic peptides, 11,728 entries for the L-form of amino acids, 229 for the D-form, and 1,215 entries for mixed forms of amino acids. Chirality information is not available for 24 peptides. We have also gathered information on peptides that have undergone chemical modifications, including C-terminal and N-terminal modifications. The chemical modifications of amino acids comprise of Dab = 2,4-diaminobutyric acid, Cha=β-cyclohexylalanine and modified terminals (2-(6-Methoxy-2-Naphthyl) propionic Acid, Leucinol (Lol, -CH(NH)-CH OH), etc. The database has 1935 entries that describe hemolytic peptides with these kinds of chemical modifications.
2. **Statistics of Natural Hemolytic Peptides**: This section presents detailed statistics on natural hemolytic peptides, including their sources, chemical modifications, functions, conformations, and stereochemistry. The database contains a total of 10,668 entries representing natural hemolytic peptides. Among these, the majority (7,557 peptides) are derived from human RBCs, followed by 737 from sheep, 711 from horse, and 467 from mouse RBCs. The remaining entries are sourced from other organisms. In terms of stereochemistry, 10,049 sequences consist exclusively of L-amino acids, 118 consist of D-amino acids, and 500 sequences are of mixed amino acid type.

**Figure 2:**
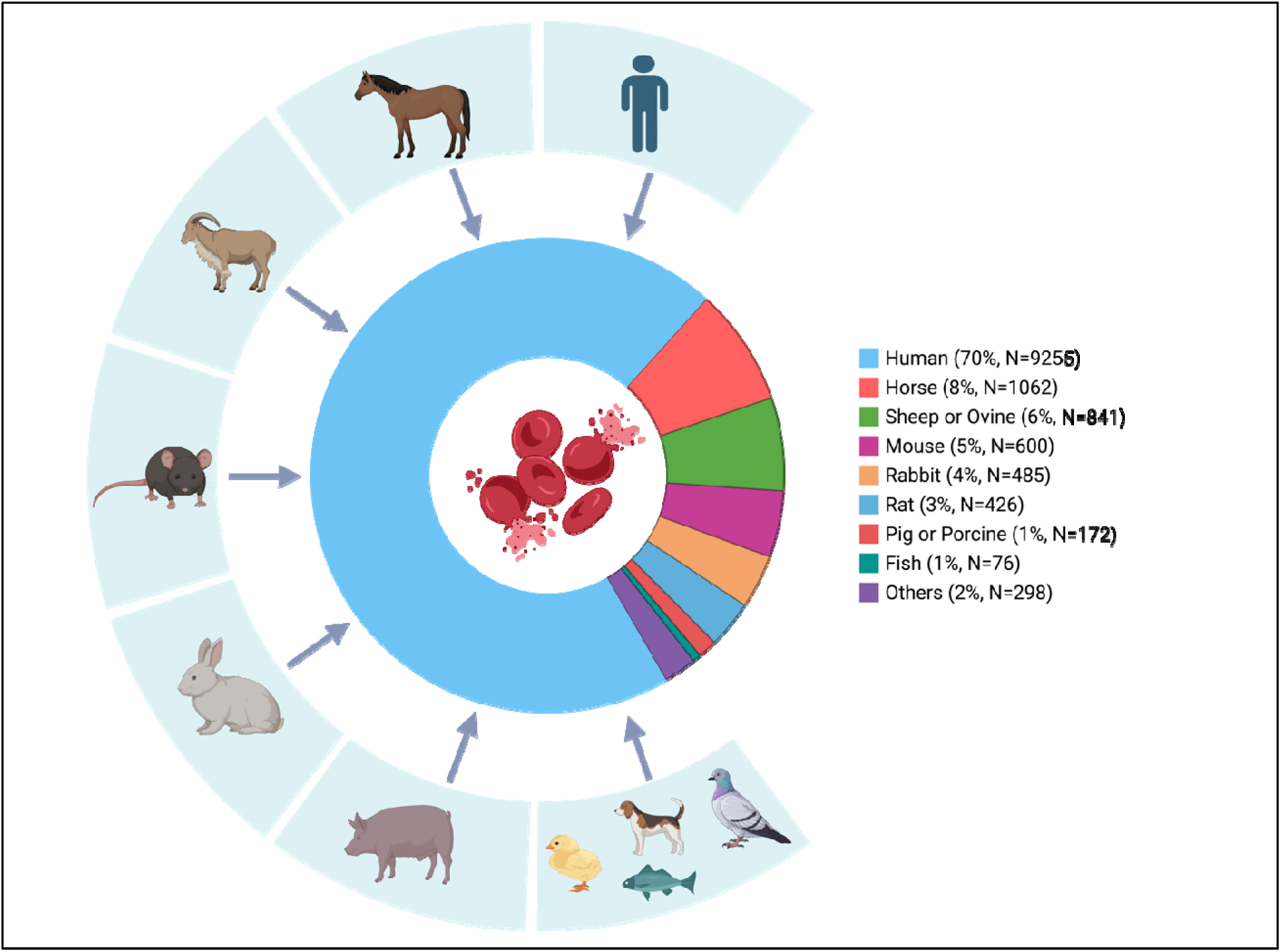
Classification of peptides based on the source of red blood cells used to assess their hemolytic activity.

### Web utility synchronization

Hemolytik 2.0 now has various easy-to-use tools for peptide analysis. The primary tools of the database are as follows:

### Search

1. **Basic search:** This variant of the search tool allows basic amenity to fetch information from the database. It enables users to perform keyword searches on various fields of the database like PMID, C-ter and N-ter modifications, stereochemistry, nature, source, hemolytic activity, origin, etc.
2. **Advanced search:** This variant of the search tool allows advanced search for the user to add multiple queries at a time and retrieve data from the database.
3. **Peptide search:** In this search tool, the user can search a given peptide against the database; there are two options: (a) Exact peptide Search-where the user can fetch information about the identical peptides in the database. (b) Containing peptide Search-it provides information based on the query peptide, which the user will input into the box.
4. **Smiles search:** This tool provides the facility to search the SMILES notation of a given peptide against the Hemolytik 2.0 peptide database in SMILES format. The RDkit tool was used to convert sequences to SMILES format.

### Browse

To make it simpler and easier for the user to discover content in the hemolytik2 database in an organized manner; we have made this browsing tool. Here, users can use an easy-to-navigate interface to explore fields including the source of the peptides (human, pig, horse, etc.), the peptide type (linear, cyclic), the stereochemistry of peptides (L, D or mix), and the non-hemolytic nature of peptides. Here, not only can the user access the fields (PMID, C-ter, and N-ter modifications, etc.) but also discover the count of each of them. Further, to make it more informative each of the fields is linked with the database so that one can get knowledge about the specific field.

### Analysis tools

1. **BLAST:** This tool allows users to perform similarity-based searches against hemolytic and non-hemolytic peptides, allowing submission of FASTA format peptide sequences and adjusting parameters like weight matrix and expectation value.
2. **Smith Waterman:** This tool conducts a similarity search more effectively for short peptides. Users can perform this analysis by inputting multiple sequences in FASTA format.
3. **Mapping:** With the tool’s sub-search and super-search features, users can map hemolytic peptides onto their peptide sequences. It extracts similar peptides based on a protein sequence, compares input peptides to all those in the Hemolytik2 database, and lets users submit sequences to find identical regions.
4. **Structure alignment:** User inputs a PDB file in the given box, and aligns its structure with the peptide structure whose ID is provided in the box.

### Download

The complete dataset from Haemolytic 2.0 is available for download. Users can access peptide sequences in multi-FASTA format, including both native and modified forms. Additionally, predicted tertiary structures are provided as PDB files. The database also supports exporting search results in CSV or Excel format for ease of analysis. All open-access reference articles from PubMed used in the curation of this resource are available for download as well.

### REST API

To facilitate automated data access, Haemolytic 2.0 now includes a RESTful API. This allows users to programmatically retrieve data based on parameters such as sequence type, peptide origin, or hemolytic activity. The API delivers responses in JSON format, enabling seamless integration with various bioinformatics pipelines and tools. Users can also parse and filter the data according to their specific requirements.

## Result and Discussion

Hemolytik 2.0, the enhanced and updated version of the Hemolytik database, now comprises a total of 13215 curated entries of hemolytic and non-hemolytic peptides. These entries were systematically compiled from an extensive review of scientific literature and multiple peptide-related databases. Each entry in the database contains comprehensive annotations, including sequence information, biological function, source organism, hemolytic activity, and structural features.

A large proportion of these peptides exhibit additional bioactivities, demonstrating the multifunctional nature of many hemolytic peptides. Specifically, Hemolytik 2.0 includes 6670 peptides with antimicrobial properties, among which 4237 exhibit antibacterial activity, 1198 are antifungal, and 697 possess antibiofilm capabilities. Additionally, the dataset features 456 peptides with anticancer activity and 196 cytotoxic peptides. Notably, 496 entries either lack functional annotations or have unknown activity profiles, highlighting potential avenues for further experimental validation.

The peptides in the database originate from a variety of species, with human-derived red blood cells (RBCs) accounting for the majority (9,255 entries). Other prominent sources include horse (1,062), sheep (841), mouse (600), rabbit (485), and rat (426) (Figure 2). An additional 298 peptides were derived from RBCs of various other organisms, including pig, porcine, and chicken, among others.

Recognizing the importance of peptide stability in therapeutic applications, Hemolytik 2.0 includes extensive information on chemically modified peptides. A total of 8340 peptides are reported to possess one or more chemical modifications aimed at enhancing their bioavailability, proteolytic resistance, or membrane permeability. Among these, C-terminal amidation is the most common modification, found in 7189 peptides. N-terminal acetylation is observed in 526 peptides, and 482 peptides feature both C-terminal amidation and N-terminal acetylation. The remaining entries contain diverse modifications, including the incorporation of D-amino acids, lipidation, and glycosylation, reflecting the growing trend of using chemical strategies to improve peptide drug performance.

One of the distinguishing features of Hemolytik 2.0 is its provision of structural information for a large number of peptides. Since the biological activity of hemolytic peptides is closely linked to their structural properties, particularly the presence of well-defined secondary structures and amphipathic organization, access to reliable structural data is crucial. Tertiary structure prediction was performed using the PEPstrMOD tool, which is tailored for modeling small peptides. Using this approach, we successfully predicted structures for 10615 peptides. These predicted models include insights into secondary structural elements (e.g., helices, sheets, coils), which are vital for understanding peptide–membrane interactions and mechanisms of hemolysis.

However, structure prediction was not feasible for a subset of chemically modified peptides due to limitations in the availability of forcefield parameters in PEPstrMOD. These peptides typically feature non-canonical amino acids or complex chemical moieties, such as 1-amino-isobutyric acid, β-naphthylalanine, norleucine, and ornithine. The absence of standard forcefields for such residues hinders reliable structural modeling. Future versions of the database aim to address this limitation by incorporating emerging forcefield libraries that support a broader range of non-standard modifications. We have also integrated the MAP (Modification and Annotation in Proteins)^20^ format, which enables the inclusion of tags directly within sequences to represent residue-level modifications such as chemical alterations, non-standard amino acids, binding sites, and mutations.

In summary, Hemolytik 2.0 not only expands the peptide dataset significantly but also enhances its utility by integrating functional, structural, and chemical information. This comprehensive resource is expected to aid researchers in elucidating structure-activity relationships and in designing novel peptides with optimized hemolytic or therapeutic potential.

The discovery of therapeutic peptides has gained attention in recent years, and its landscape continues to evolve at an impressive pace, driven by significant technological breakthroughs and a burgeoning body of research^1^. Hemolytic toxicity is a problem for many promising peptides, because it may cause red blood cells to be accidentally destroyed, which would restrict their therapeutic usage and raise serious safety issues when used in clinical use. The emergence and ongoing refinement of comprehensive hemolytic peptide databases represent a pivotal turning point in addressing this critical issue. These tools provide a consolidated platform for comprehending the complex link between peptide sequence, structure, and hemolytic activity, beyond the constraints of empirical screening and scattered information^21^. The first extensive database of its sort, Hemolytik, offers experimental data on hemolytic peptides and their potencies.

By providing curated experimental data, these databases empower researchers to move towards a rational design paradigm^22^. They serve as the bedrock for developing increasingly accurate *in silico* prediction tools, enabling early identification of potentially toxic candidates and facilitating virtual screening of vast peptide libraries. Ultimately, the significance of hemolytic peptide databases lies in their ability to accelerate the development of safer and more translatable peptide therapeutics^23^. By mitigating a major obstacle in the drug development pipeline, these resources pave the way for a future where the unique advantages of peptides – their specificity, potency, and potential for targeted delivery^24^ can be fully realized for the benefit of human health. The continued growth, integration with other biological data^25,26^, and accessibility of these databases are not merely desirable, but essential for unlocking the full therapeutic potential of peptides and ensuring their successful journey from bench to bedside^23,27^.

### Comparison with the Previous Version

Hemolytik 2.0 represents a substantial advancement over its predecessor in both data content and annotation quality. The data sources and literature coverage have also expanded significantly. While the original database used APD2, CAMP, UniProt, and DAMPD, Hemolytik 2.0 incorporates updated and broader resources such as APD3, CAMPR4, UniProt, and DRAMP 4.0. The number of PubMed articles referred for initial screening has increased from approximately 900 to 4,350, and the number of publications manually curated for extracting peptide information has risen from 446 to 1,228. The initial version of Hemolytik comprised 2970 peptide entries, whereas Hemolytik 2.0 includes 13215 curated entries, signifying more than a fourfold increase. The number of unique hemolytic peptides has grown from 1750 to 5598, and non-hemolytic peptides have expanded from 295 to 2410 entries (Figure 3). Moreover, the variety of red blood cell (RBC) sources used in hemolysis assays has also increased, from 17 species in the earlier version to 20 in the updated release.

**Figure 3:**
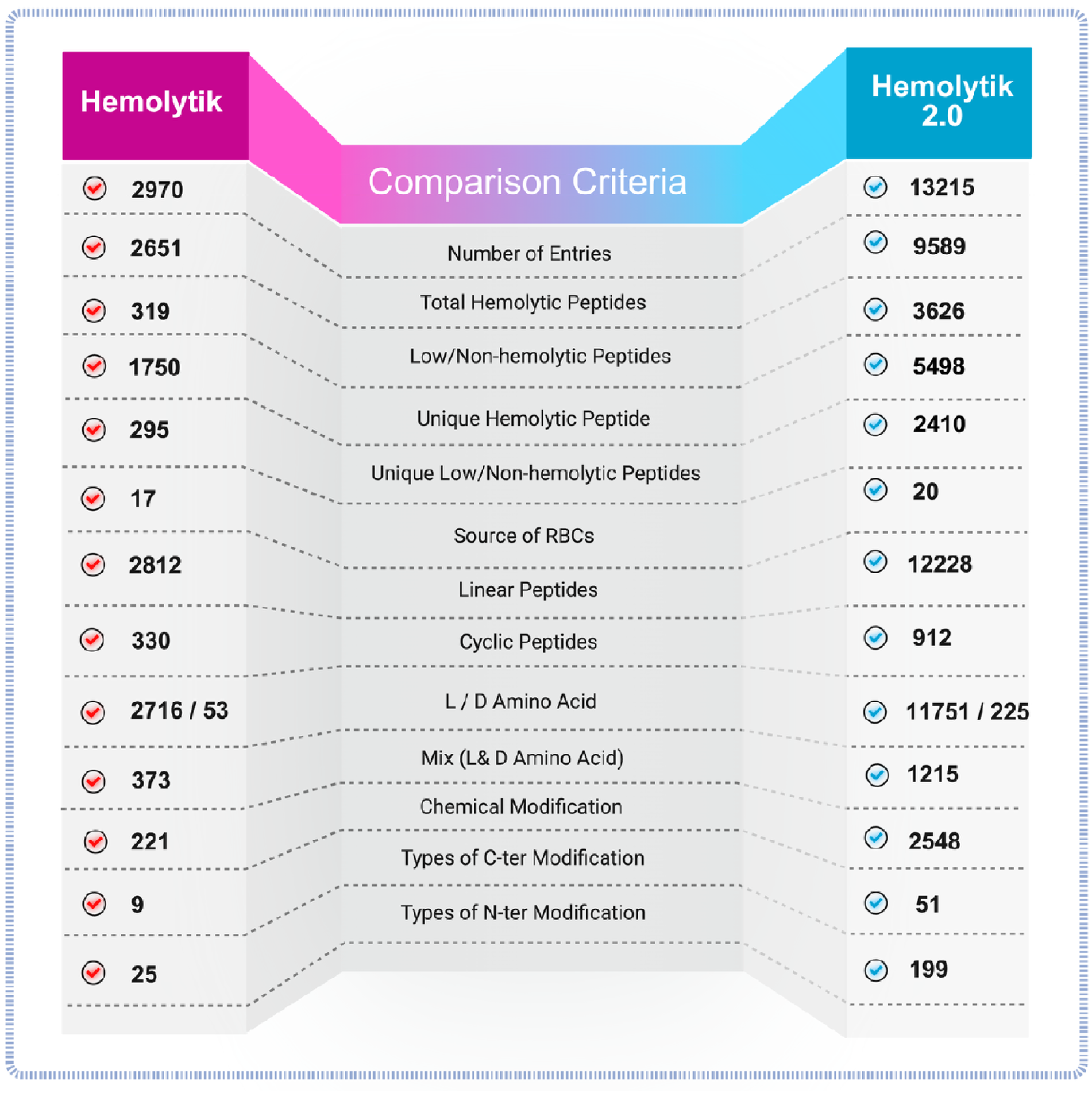
A visual summary highlighting the advancements in data volume, diversity, and structural information in Hemolytik 2.0 compared to its previous version.

Structurally, Hemolytik 2.0 demonstrates a more comprehensive classification of peptide topologies. While the first version primarily focused on linear (2669) and cyclic (301) peptides, the updated version includes 12228 linear peptides, 912 cyclic peptides, and 75 peptides with complex topologies such as branched, bicyclic, stapled, and macrocyclized architectures. Additionally, Hemolytik 2.0 offers a broader representation of peptide stereochemistry. The number of entries composed exclusively of L-amino acids has increased from 2,583 to 11751, D-amino acids from 47 to 225, and mixed stereochemistry peptides from 373 to 1215.

Chemical modifications are better represented in the current version, with the number of chemically modified peptides growing from 221 to 2548. A significant improvement lies in the detailed annotation of terminal modifications: the earlier version provided limited information on 9 C-terminal and 25 N-terminal modifications, while Hemolytik 2.0 now documents 51 types of C-terminal and 199 types of N-terminal modifications, including amidation, acetylation, lipidation, and other structural enhancements. Collectively, these improvements position Hemolytik 2.0 as a significantly enriched and more versatile platform for studying hemolytic peptides, providing an essential resource for researchers in peptide therapeutics, drug development, and membrane biology.

### Limitations and Updates of Hemolytik 2.0

Hemolytik 2.0 provides extensive structural and functional data for hemolytic and non-hemolytic peptides; however, it has certain limitations. One significant challenge lies in its inability to predict the structures of chemically modified peptides. The presence of non-standard amino acids (e.g., Bip = 4,4′-biphenyl-L-alanine, Cha = cyclohexylalanine, Dab = diaminobutyric acid, Ornithine) and terminal modifications (e.g., stearoyl, Valinol) often leads to structural complexities that are not well-represented in conventional structural databases or modeling tools. Consequently, accurate prediction of SMILES notations or DSSP-based secondary structures for these peptides is currently hindered. However, with the progressive development of force field libraries encompassing modified residues, future updates may enable reliable structural prediction for such peptides.

To ensure continuous improvement, Hemolytik 2.0 offers an HTML-based submission interface, allowing users to contribute newly identified hemolytic and non-hemolytic peptides backed by experimental validation. Each submission is manually reviewed and verified for authenticity prior to inclusion, maintaining the database’s integrity and reliability. The database is regularly updated by our team to incorporate the latest research findings and user-contributed data, ensuring it remains a comprehensive and up-to-date resource for the scientific community.

## Availability

The database Hemolytik2.0 is freely accessible at https://webs.iiitd.edu.in/raghava/hemolytik2/ and data can be downloaded from the downloads tab given in the website or directly from API at https://webs.iiitd.edu.in/raghava/hemolytik2/api/rest.html by selecting the query fields.

## Authors Contribution

AS, KSA, and ASR manually collected the data. AS, KSA, and ASR. manually curated and analyzed the data. KSA and AS developed the backend and front end of the web server. ASR and GPSR prepared the manuscript. ASR, KSA, AS, and GPSR reviewed the manuscript. GPSR conceived and coordinated the project. All authors read and approved the final manuscript.

## Acknowledgments

Authors are thankful to the All India Council for Technical Education (AICTE), University Grants Commission (UGC), Indraprastha Institute of Information Technology (IIITD) for fellowships and financial support, and the Department of Computational Biology, IIITD, New Delhi, for infrastructure and facilities. We thank the DBT for providing an infrastructure grant to the institute. Furthermore, they would like to acknowledge BioRender.com for creating the figures utilized in this work.

## References

1. Wang, L., Wang, N., Zhang, W., Cheng, X., Yan, Z., Shao, G., Wang, X., Wang, R., & Fu, C. (2022) Therapeutic peptides: current applications and future directions. Signal Transduction and Targeted Therapy [Internet] 7,1 48. Available from: https://www.nature.com/articles/s41392-022-00904-4

2. Rossino G, Marchese E, Galli G, Verde F, Finizio M, Serra M, Linciano P, Collina S (2023) Peptides as therapeutic agents: Challenges and opportunities in the Green transition era. Molecules [Internet] 28. Available from: 10.3390/molecules28207165

3. Mullard A (2024) 2023 FDA approvals. Nat. Rev. Drug Discov. [Internet] 23:88–95. Available from: 10.1038/d41573-024-00001-x

4. Al Musaimi O, Al Shaer D, Albericio F, de la Torre BG (2023) 2022 FDA TIDES (peptides and oligonucleotides) harvest. Pharmaceuticals (Basel) [Internet] 16. Available from: 10.3390/ph16030336

5. Al Shaer D, Al Musaimi O, Albericio F, de la Torre BG (2024) 2023 FDA TIDES (peptides and oligonucleotides) harvest. Pharmaceuticals (Basel) [Internet] 17:243. Available from: 10.3390/ph17020243

6. Rathore AS, Jain S, Choudhury S, Raghava GPS (2025) NTxPred2: A large language model for predicting neurotoxic peptides and neurotoxins. bioRxiv [Internet]. Available from: 10.1101/2025.03.01.640936

7. Rathore AS, Choudhury S, Arora A, Tijare P, Raghava GPS (2024) ToxinPred 3.0: An improved method for predicting the toxicity of peptides. Comput. Biol. Med. [Internet] 179:108926. Available from: 10.1016/j.compbiomed.2024.108926

8. Sharma N, Naorem LD, Jain S, Raghava GPS (2022) ToxinPred2: an improved method for predicting toxicity of proteins. Brief. Bioinform. [Internet] 23. Available from: 10.1093/bib/bbac174

9. Rathore AS, Kumar N, Choudhury S, Mehta NK, Raghava GPS (2025) Prediction of hemolytic peptides and their hemolytic concentration. Commun. Biol. [Internet] 8:176. Available from: 10.1038/s42003-025-07615-w

10. Tosteson MT, Holmes SJ, Razin M, Tosteson DC (1985) Melittin lysis of red cells. J. Membr. Biol. [Internet] 87:35–44. Available from: 10.1007/bf01870697

11. Chen L, Patrone N, Liang JF (2012) Peptide self-assembly on cell membranes to induce cell lysis. Biomacromolecules [Internet] 13:3327–3333. Available from: 10.1021/bm301106p

12. Spaller BL, Trieu JM, Almeida PF (2013) Hemolytic activity of membrane-active peptides correlates with the thermodynamics of binding to 1-palmitoyl-2-oleoyl-sn-glycero-3-phosphocholine bilayers. J. Membr. Biol. [Internet] 246:257–262. Available from: 10.1007/s00232-013-9525-z

13. Chaudhary K, Kumar R, Singh S, Tuknait A, Gautam A, Mathur D, Anand P, Varshney GC, Raghava GPS (2016) A web server and mobile app for computing hemolytic potency of peptides. Sci. Rep. [Internet] 6. Available from: 10.1038/srep22843

14. Timmons PB, Hewage CM (2020) HAPPENN is a novel tool for hemolytic activity prediction for therapeutic peptides which employs neural networks. Sci. Rep. [Internet] 10. Available from: 10.1038/s41598-020-67701-3

15. Kumar V, Kumar R, Agrawal P, Patiyal S, Raghava GPS (2020) A method for predicting hemolytic potency of chemically modified peptides from its structure. Front. Pharmacol. [Internet] 11:54. Available from: 10.3389/fphar.2020.00054

16. Wang G, Li X, Wang Z (2016) APD3: the antimicrobial peptide database as a tool for research and education. Nucleic Acids Res. [Internet] 44:D1087–93. Available from: 10.1093/nar/gkv1278

17. UniProt Consortium (2023) UniProt: The universal protein knowledgebase in 2023. Nucleic Acids Res. [Internet] 51:D523–D531. Available from: 10.1093/nar/gkac1052

18. Gawde U, Chakraborty S, Waghu FH, Barai RS, Khanderkar A, Indraguru R, Shirsat T, Idicula-Thomas S (2023) CAMPR4: a database of natural and synthetic antimicrobial peptides. Nucleic Acids Res. [Internet] 51:D377–D383. Available from: 10.1093/nar/gkac933

19. Seshadri Sundararajan V, Gabere MN, Pretorius A, Adam S, Christoffels A, Lehväslaiho M, Archer JAC, Bajic VB (2012) DAMPD: a manually curated antimicrobial peptide database. Nucleic Acids Res. [Internet] 40:D1108–12. Available from: 10.1093/nar/gkr1063

20. Shendre A, Mehta NK, Rathore AS, Kumar N, Patiyal S, Raghava GPS (2025) MAP format for representing chemical modifications, annotations, and mutations in protein sequences: An extension of the FASTA format. arXiv [q-bio.BM] [Internet]. Available from: 10.48550/ARXIV.2505.03403

21. Gautam A, Chaudhary K, Singh S, Joshi A, Anand P, Tuknait A, Mathur D, Varshney GC, Raghava GPS (2014) Hemolytik: a database of experimentally determined hemolytic and non-hemolytic peptides. Nucleic Acids Res. [Internet] 42:D444–9. Available from: 10.1093/nar/gkt1008

22. Skowron KJ, Baliga C, Johnson T, Kremiller KM, Castroverde A, Dean TT, Allen AC, Lopez-Hernandez AM, Aleksandrova EV, Klepacki D, et al. (2023) Structure-activity relationships of the antimicrobial peptide natural product apidaecin. J. Med. Chem. [Internet] 66:11831–11842. Available from: 10.1021/acs.jmedchem.3c00406

23. Fosgerau K, Hoffmann T (2015) Peptide therapeutics: current status and future directions. Drug Discov. Today [Internet] 20:122–128. Available from: 10.1016/j.drudis.2014.10.003

24. Ioannou P, Baliou S, Kofteridis DP (2023) Antimicrobial peptides in infectious diseases and beyond-A narrative review. Life (Basel) [Internet] 13. Available from: 10.3390/life13081651

25. Seelig J (2004) Thermodynamics of lipid-peptide interactions. Biochim. Biophys. Acta [Internet] 1666:40–50. Available from: 10.1016/j.bbamem.2004.08.004

26. Matsuzaki K (1999) Why and how are peptide-lipid interactions utilized for self-defense? Magainins and tachyplesins as archetypes. Biochim. Biophys. Acta [Internet] 1462:1–10. Available from: 10.1016/s0005-2736(99)00197-2

27. Craik DJ, Fairlie DP, Liras S, Price D (2013) The future of peptide-based drugs. Chem. Biol. Drug Des. [Internet] 81:136–147. Available from: 10.1111/cbdd.12055

